# Mass spectrometry imaging links interxylary cork formation with phytochemical distribution in *Scutellaria baicalensis*

**DOI:** 10.1101/2023.11.17.567626

**Authors:** Lieyan Huang, Lixing Nie, Jing Dong, Lingwen Yao, Shuai Kang, Hongyu Jin, Feng Wei, Shuangcheng Ma

## Abstract

Interxylary cork, an anomalous cork structure of plant, is occasionally seen in older roots of *Scutellaria baicalensis* Georgi (Lamiaceae). Xylem tissues encircled by interxylary corks gradually decay. Efforts have been made to elucidate the development of interxylary cork in *S. baicalensis*, but the variation of phytochemical accumulation in different root tissues segmented by interxylary cork has not been studied. In this investigation, mass spectrometry imaging was employed to visualize the *in situ* phytochemicals in the decayed root of *S. baicalensis* (Kuqin). A special flavonoid was observed to show especially high mass intensity in the cork region, and its spatial distribution was regarded as a reflection of cork development. Interxylary corks were found to be successively formed in the root of *S. baicalensis*. After the continuous formation of interxylary corks, root tissues were divided into different parts by interxylary corks, including the non-decayed part, slightly decayed part and severely decayed part. Interestingly, pharmaceutically important flavonoids presented different accumulation tendencies in tissues of different decaying degrees. Owing to the non-targeted analytical function of mass spectrometry imaging, the holistic difference of chemical composition of decayed and non-decayed tissues of Kuqin was deciphered. To our knowledge, this is the first experiment that maps out the successive formation of interxylary corks in *S. baicalensis* using mass spectrometry imaging. Also, our research successfully correlates plant anatomy with the spatial distribution of phytochemicals, which sets a good example for the exploration of plant science from the dimension of spatial information.

## Introduction

Scutellariae Radix, the dried root of *Scutellaria baicalensis* Georgi, has been widely applied to treat diseases such as hepatitis, diarrhoea, inflammation and respiratory infections (Zhao *et al*., 2019). Traditionally, there are two classifications of commercial specifications of Scutellariae Radix, namely, Kuqin and Ziqin. These two kinds of medicinal materials are collected at different growing stages of *S. baicalensis*. As the plant grows into older ages, some of the roots start to decay from the inside. The decayed pith is separated from normal tissues by the development of interxylary cork. The older root having a decayed pith is called Kuqin, while the younger root having a solid texture is called Ziqin. Though Ziqin and Kuqin have the same botanical origin, they are believed to exhibit different pharmacological effects. It was proved by a modern study that Ziqin showed better therapeutic effect when treating large intestine diseases (Zhang *et al*., 2020). Another investigation stated that Kuqin had better anti-inflammatory and analgesic activities than Ziqin (Sun *et al*., 2023a). The chemical difference of Kuqin and Ziqin should be the inherent reason why two kinds of Scutellariae Radix hold different pharmacological effects. The formation of a decayed pith is the vital step contributing to the transformation from Ziqin into Kuqin. As a result, it is of great importance to dig out the chemical difference of decayed and non-decayed tissues of Kuqin.

Before chemical analysis, the anatomical structure of Kuqin should be illustrated first. Interxylary cork, which serves as a boundary between abnormal xylem and healthy xylem in decayed *S. baicalensis*, has gained the attention of researchers. Interxylary cork is an anomalous status of cork (phellem) that originates from the inner tissues of secondary xylem. Early in 1934, Moss identified the existence of interxylary cork in a massive number of plants (Moss, 1934). Similar to general cork that develops along the outer edge of stems, roots, tubers, fruit and healing tissues (Waisel, 1995; Bernal *et al*., 2008), interxylary cork also acts as a protective tissue that prevents plant from excessive water loss, pathogen attack and ultraviolet irradiation (Lendzian, 2006). As was proposed by Moss and Gorham in 1953, the development of interxylary cork helped plants to resist against desiccation and adapt to their habitats, mainly dry and wind-blown soil (Moss & Gorham, 1953). The main habitats of *S. baicalensis* in China is Hebei and Neimenggu, where drought and strong wind frequently take place. Therefore, the development of interxylary cork in *S. baicalensis* can be attributed to the geographical characteristics of these habitats.

The formation of interxylary cork in *S. baicalenis* has been recently revealed using optic microscopic and electron microscopic (Wang *et al*., 2020). It was stated that tylosis or other substances occurring in some vessels were the trigger of interxylary cork formation. Parenchyma cells adjacent to these abnormal vessels recovered meristematic abilities and differentiated into new cork cambium (phellogen). The activity of meristematic cambium cells produced more cell layers both inwardly and outwardly, forming phelloderm and interxylary cork. Later on, a mature interxylary cork in the shape of a large ring was formed. Ultimately, the tissues enclosed by a mature interxylary cork gradually rot or shed, leading to the formation of a decayed pith in Kuqin (Wang *et al*., 2020). Interestingly, two other papers reported the development of interxylary cork as a multicentric process in *Gentiana macrophylla* Pall. and *Astragalus membranaceus* (Fisch) Bge. var. *mongholicus* (Bge.) Hsiao (Luo, 1987; Han *et al*., 2021). Small interxylary corks could be successively or simultaneously formed at different positions of xylem, and the integration of which contributed to the expansion of interxylary cork. However, the sequential formation of multiple interxylary corks has not been mentioned in the anatomical study of *S. baicalensis*, which arouses question on the development of interxylary cork in *S. baicalensis*.

Currently, region-specific analysis of Kuqin is usually carried out using liquid chromatography (LC) or liquid chromatography coupled with mass spectrometry (LC-MS) (Wang *et al*., 2012; Song *et al*., 2020). The HPLC fingerprint chromatograms of decayed and non-decayed tissues of Kuqin were found to differ a lot from each other (Wang *et al*., 2012). Another investigation illustrated the varied contents of bioactive compounds in decayed and non-decayed regions of Kuqin, but this analysis was limited to several important flavonoids (Song *et al*., 2020). During LC or LC-MS analysis, the homogenization of sample materials was an unavoidable step, which led to the loss of spatial information of detected compounds (Tang *et al*., 2019). Besides, the extraction of natural compounds from dissected tissues might cause the degradation of target compounds (Nie *et al*., 2021). As a result, a technique which allows untargeted detection of natural compounds in native tissues of Kuqin is of urgent importance.

Different from classical analytical tools, mass spectrometry imaging (MSI) requires no sample homogenization, thus is able to maintain the spatial information of detected compounds during analytical tasks (Huang *et al*., 2022). Meanwhile, MSI supports the simultaneous detection of numerous classes of natural compounds (Bjarnholt *et al*., 2014). Owing to its superiorities, MSI has been widely adopted to uncover the biosynthesis, transportation and accumulation of natural compounds in plants (Husain *et al*., 2020; Sun *et al*., 2023b). Nevertheless, seldom has been done to decipher the development of plant structure using MSI, not to mention the linkage between phytochemical distribution and plant anatomy.

In this paper, atmospheric pressure matrix assisted laser desorption and ionization-quadrupole-time of flight-mass spectrometry imaging (AP-MALDI-Q-TOF-MSI) was employed to assist in the exploration of plant anatomy and phytochemistry of *S. baicalensis*. The continuous development of interxyalry corks in *S. baicalensis* was mapped out by the ion image of skullcapflavone Ⅱ. The spatial distribution of natural compounds was found to be related with the decaying stages of different tissues. Holistic chemical composition of decayed and non-decayed tissues of *S. baicalensis* was compared via multivariate analysis. Overall, this investigation succeeded to link phytochemical distribution with plant structure, providing new inspirations for the study on anatomy and phytochemistry of medicinal plants.

## Results

### Spatial distribution of skullcapflavone Ⅱ was discovered as a reflection of interxylary cork development

With the employment of MSI, a collection of natural compounds was simultaneously detected within a single run. The *in situ* distribution of phytochemicals in Kuqin was intuitively visualized in the mass spectrometry (MS) images. In our previous work, a few chemicals were found to exhibit especially high mass intensities in the root cork of *S. baicalensis*, including the compound detected at *m/z* 413.0631, which was putatively identified as skullcapflavone Ⅱ (Huang *et al*., 2023). In this investigation, high mass intensity of skullcapflavone Ⅱ was also observed in the interxylary cork of roots (Fig. 1B), indicating that interxylary cork was a preferential accumulation site of skullcapflavone Ⅱ. Therefore, the spatial distribution of skullcapflavone Ⅱ could be regarded as a reflection of interxylary cork development.

**Fig. 1.**
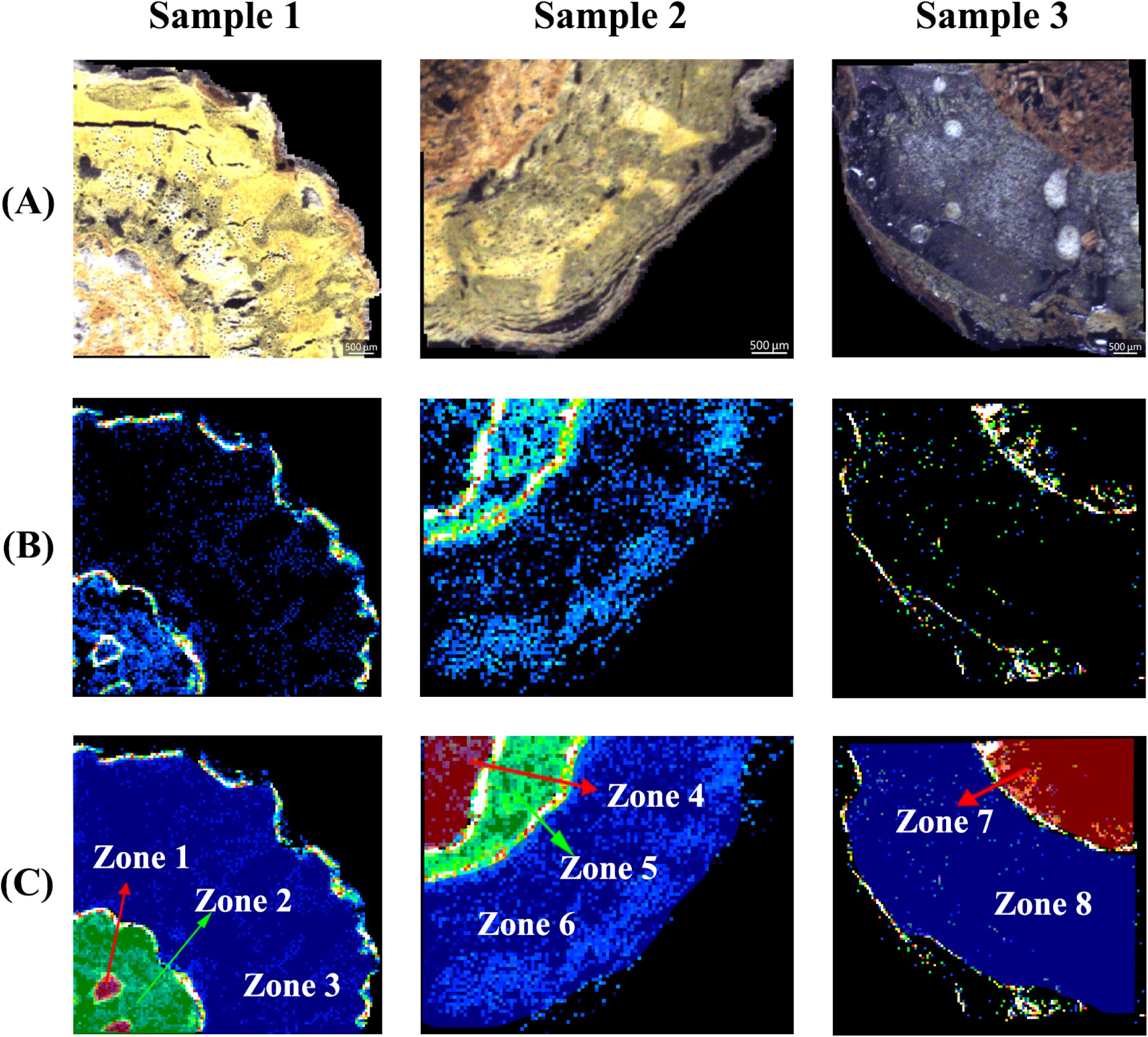
Optical images and MS images generated from the transverse sections of three Kuqin samples. (A) Optical images of the transverse section of three Kuqin samples. The optical images were captured by the charge coupled device camera attached to the iMScope QT instrument. (B) MS images of skullcapflavone Ⅱ (*m/z* 413.0645, [C_17_H_14_O_8_+K]^+^, Δ=1.45 ppm). The spatial distribution of skullcapflavone Ⅱ was found to be related to the development of interxylary cork. (C) Segmentation of root tissues according to the development of interxylary corks. The root sections were divided into different zones, the tissue within each zone was assumed to present a different decaying degree compared with other zones.

The decayed center of Kuqin was separated from healthy tissues by the development of interxylary cork (Wang *et al*., 2020). By comparing with the optical image of each sample (Fig. 1A), the MSI signal of skullcapflavone Ⅱ was observed to draw the edge of the decayed pith in three Kuqin samples (Fig. 1B). Interestingly, in sample 1 and sample 2, high mass intensigy of skullcapflavone Ⅱ was observed in the form of multiple rings (Fig. 1B). Interxylary corks in the shape of smaller rings were found to exist inside the large interxylary cork of sample 1, indicating a multicentric development of small interxylary corks in the center of Kuqin (Fig. 1B). Whereas in sample 2, two layers of interxylary corks were presented in the shape of concentric circles (Fig. 1B). The distribution of skullcapflavone Ⅱ in sample 1 and sample 2 suggested the successive development of interxylary corks in Kuqin.

In the optical images (Fig. 1A), tissues within the decayed center presented different colors, the darker the color of tissues, the severer the decaying degree of tissues. By comparing with MS images, it was obvious to find that the signal of skullcapflavone Ⅱ particularly outlined the margin of tissues of different decaying degrees. In other words, the development of interxylary corks divided the whole root tissue into different zones, each zone was of a different decaying degree compared with others (Fig. 1C).

### Bioactive flavonoids were found to show varied accumulation tendencies in tissues segmented by interxylary corks

Baicalein, wogonin, baicalin and wogonoside were the four major flavonoids in Scutellariae Radix. On-tissue tandem mass analysis of four flavonoids were employed, and the fragments of which were in consistency with those generated from reference standards. MS images of four flavonoids were constructed according to the *m/z* values of the protonated ions of four flavonoids (Fig. 2, A to D). Apparently, baicalein and wogonin exhibited different accumulation tendencies compared with their glucuronides.

**Fig. 2.**
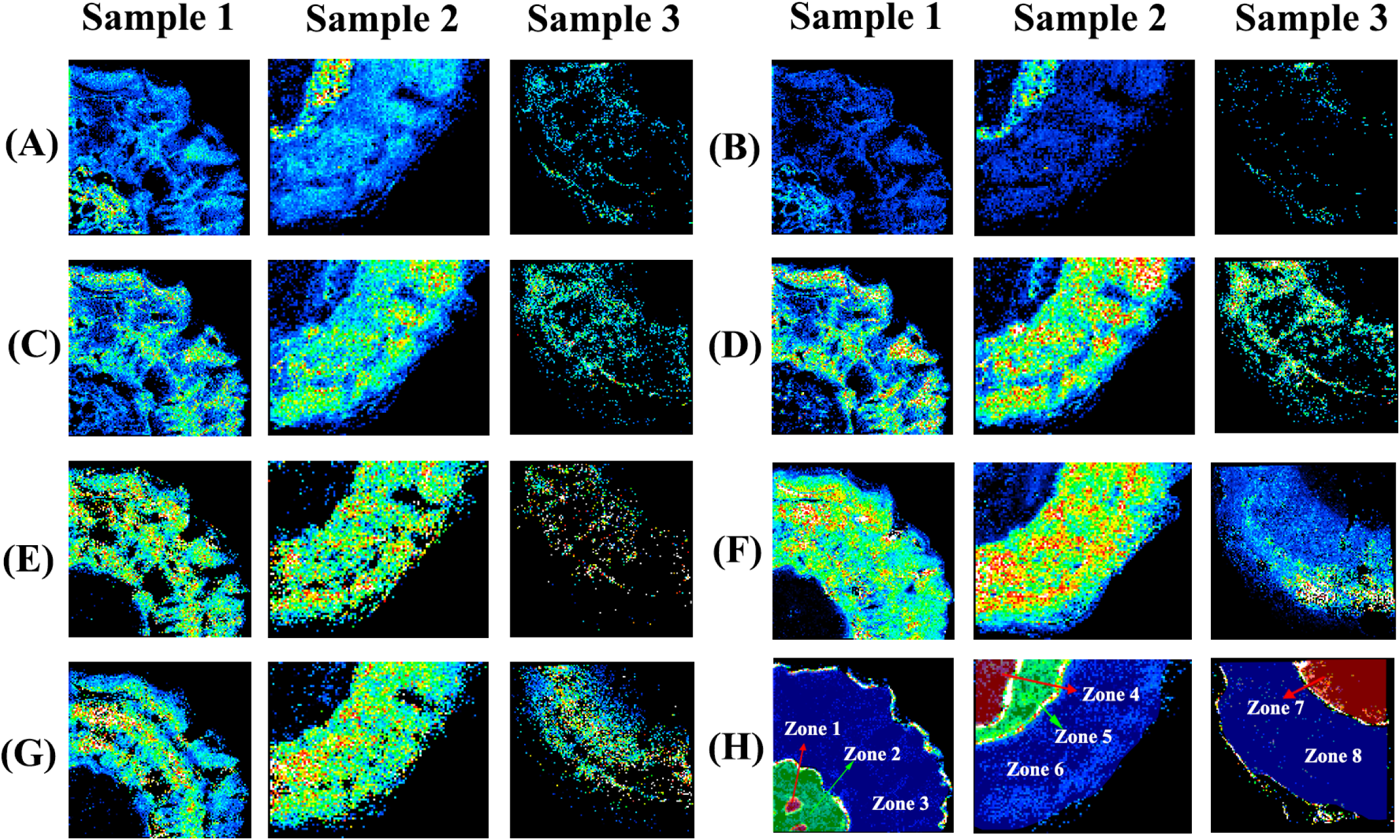
MS images of four major flavonoids and saccharides in three Kuqin samples. (A) MS images of baicalein (*m/z* 309.0160, [C_15_H_10_O_5_+K]^+^, Δ=1.62 ppm). (B) MS images of wogonin (*m/z* 323.0316, [C_16_H_12_O_5_+K]^+^, Δ=1.86 ppm). (C) MS images of baicalin (*m/z* 485.0486, [C_21_H_18_O_11_+K]^+^, Δ=0 ppm). (D) MS images of wogonoside (*m/z* 499.0637, [C_22_H_20_O_11_+K]^+^, Δ=1.20 ppm). (E) MS images of glucose (*m/z* 219.0269, [C_6_H_12_O_6_+K]^+^, Δ=0.91 ppm). (F) MS images of sucrose (*m/z* 381.0800, [C_12_H_22_O_11_+K]^+^, Δ=0.26 ppm). (G) MS images of raffinose (*m/z* 543.1323, [C_18_H_32_O_16_+K]^+^, Δ=0.92 ppm). (H) Segmentation of root tissues according to the development of interxylary corks.

Two glucuronides, baicalin and wogonoside, were found to show relatively higher mass intensity in the non-decayed part of three Kuqin samples (Fig. 2, C and D). Whereas in decayed region of Kuqin, the intensity of baicalin and wogonoside sharply decreased. It seemed that the existence of a decayed pith hindered the accumulation of baicalin and wogonoside. The distribution modes of baicalin and wogonoside were consistent with those of saccharides. Glucose, sucrose, raffinose and/or their isomers were also observed to exhibit especially high mass signals in the non-decayed part of Kuqin (Fig. 2, E to G). It could conceivably be hypothesized that after an interxylary cork was formed, the horizontal transportation of saccharides was cut off by the interxylary cork. Therefore, in decayed xylem tissue enclosed by interxylary cork, the synthesis of baicalin and wogonoside was decreased because of a lack of substrates for glycosation.

Two flavonoid aglycones, baicalein and wogonin, presented more complicated distribution modes (Fig. 2, A and B) compared with their glucuronides. As was mentioned in the previous section, successively formed interxylary corks could be utilized to divide decayed tissues into slightly decayed part and severely decayed part. In sample 1, the relative intensities of baicalein and wogonin showed higher intensities in zone 2, where xylem tissue was going through a slightly decaying process. For the severely decayed tissue (zone 1) which was enclosed by a newly formed interxylary cork, the mass intensities of baicalein and wogonin significantly dropped. Similarly, in sample 2, mass intensities of baicalein and wogonin were more intense in slightly decayed tissue (zone 5) compared with severely decayed tissue (zone 4). In sample 3, baicalein and wogonin were scarcely distributed in the decayed pith of Kuqin. The MS images of two flavonoid aglycones intuitively illustrated that the relative content of baicalein and wogonin varied between tissues segmented by interxylary corks. In other words, the spatial distribution of baicalein and wogoin was probably related to the decaying extent of tissues.

### Principal component analysis deciphered the holistic chemical difference of decayed and non-decayed xylem of Kuqin

The transformation of healthy xylem into abnormal xylem is the key point contributing to the difference of Kuqin and Ziqin. Region-specific analysis of Kuqin was readily achieved with the employment of MSI. With reference to the optical image, each longitudinal section was divided into different regions of interest (ROI). For each longitudinal section, five ROIs were drawn in decayed xylem region, and the other five ROIs were drawn in non-decayed xylem region (Fig. 3). Average mass spectra were obtained from the mass spectrometry data generated from the native tissue confined within different ROIs. From the typical mass spectra generated from ROIs drawn in decayed and non-decayed xylem (Fig. 4, A to D), it was obvious to find that the chemical composition of decayed and non-decayed xylem differed a lot from each other.

**Fig. 3.**
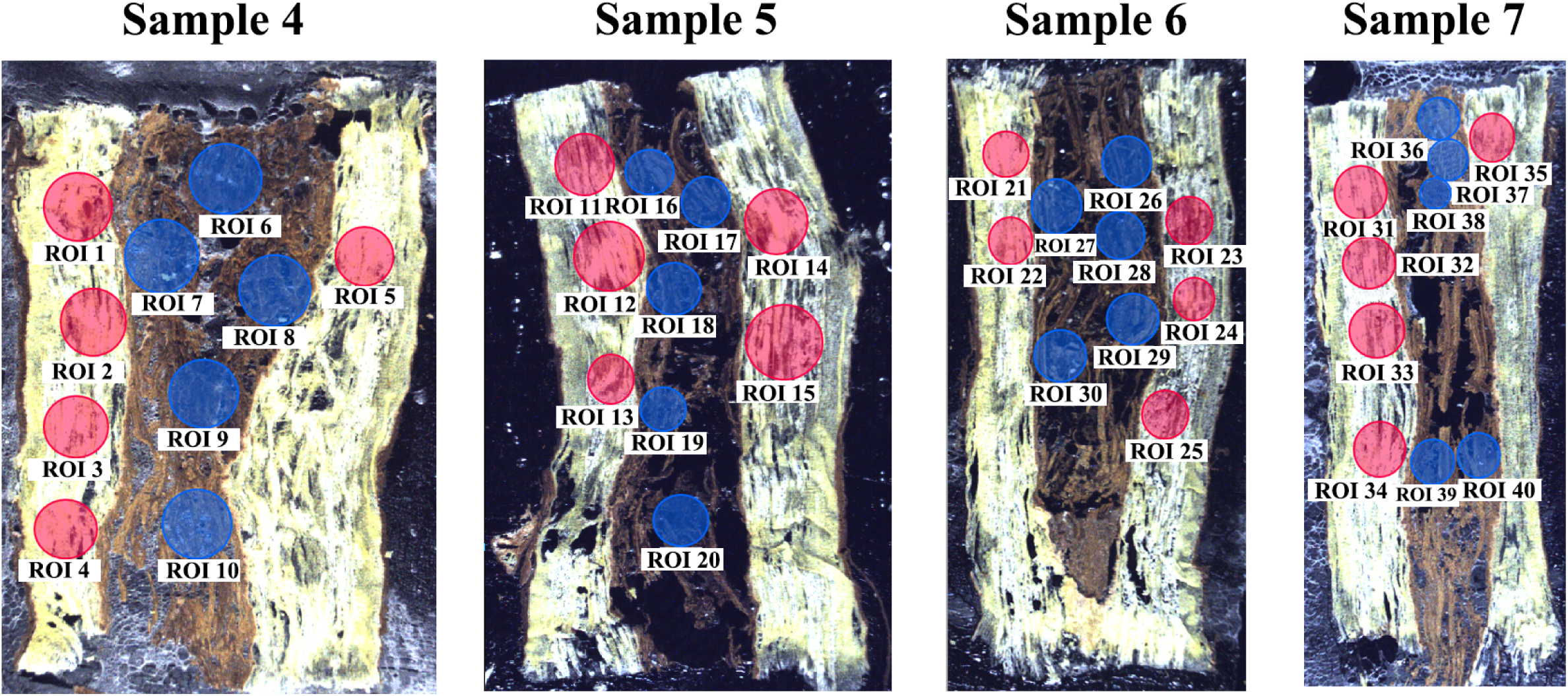
Illustration of ROI selection within the longitudinal sections of Kuqin samples. For each sample, five ROIs were randomly drawn within the decayed xylem, while the other five were drawn within the non-decayed xylem.

**Fig. 4.**
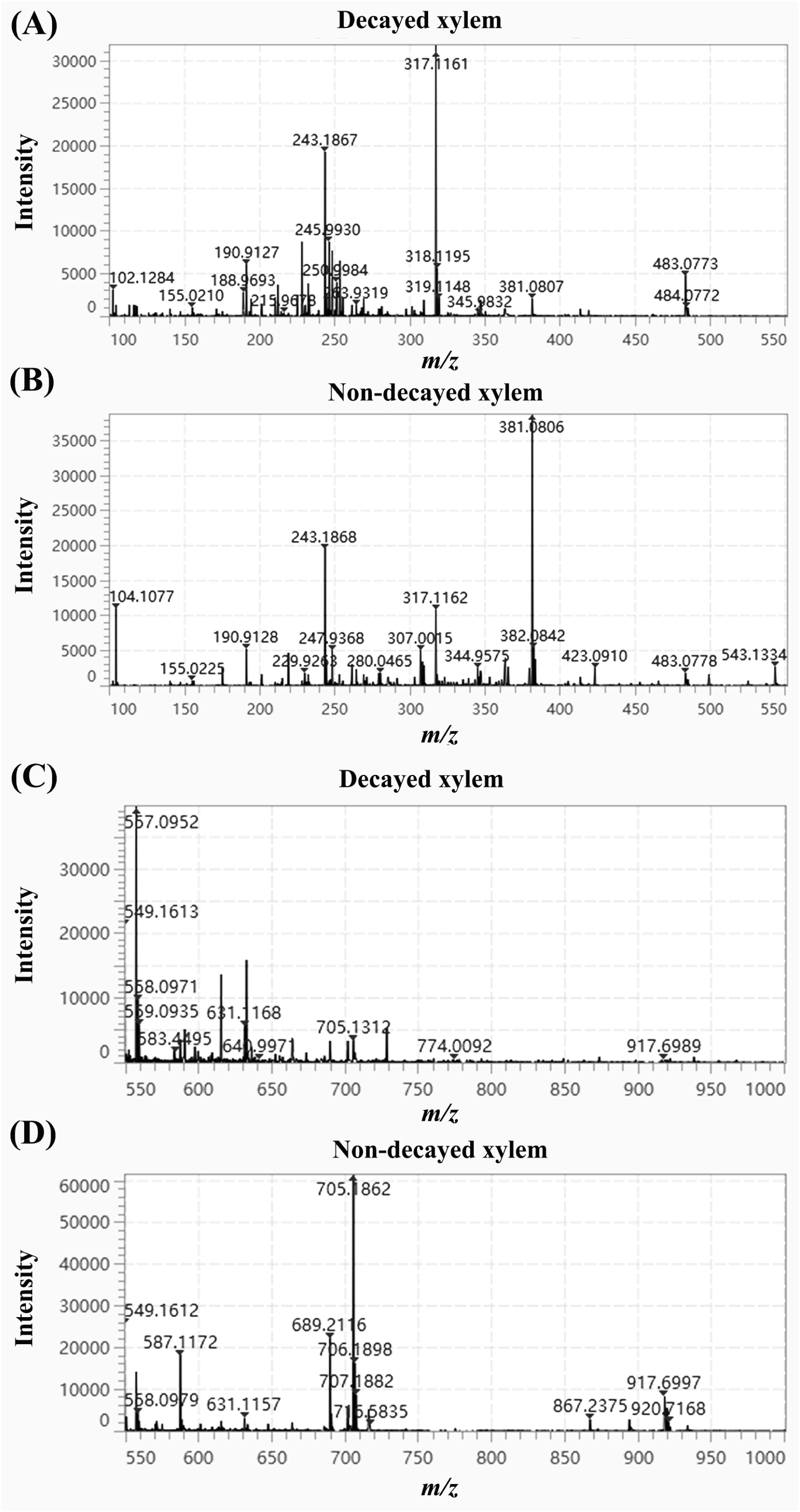
Average mass spectra generated from representative ROIs drawn in decayed and non-decayed regions of sample 4. (A) An average mass spectrum generated from ROI 6 drawn in the decayed xylem, acquisition mass range *m/z* 100-550. (B) An average mass spectrum generated from ROI 1 drawn in the non-decayed xylem, acquisition mass range *m/z* 100-550. (C) An average mass spectrum generated from ROI 6 drawn in the decayed xylem, acquisition mass range *m/z* 550-1000. (D) An average mass spectrum generated from ROI 1 drawn in the non-decayed xylem, acquisition mass range *m/z* 550-1000.

Principal component analysis (PCA) was then carried out to compare the chemical difference between decayed and non-decayed xylem of Kuqin. Unambiguous differentiation of decayed and non-decayed xylem was achieved using PCA. As illustrated in the score plots, blue dots representing decayed xylem regions were clearly separated from red dots representing non-decayed xylem regions (Fig. 5, A and B). According to the loading vector of detected chemicals, differential ions that contributed most to the difference of two groups were sorted out, and were highlighted with purple color in the loading plots (Fig. 5, A and B).

**Fig. 5.**
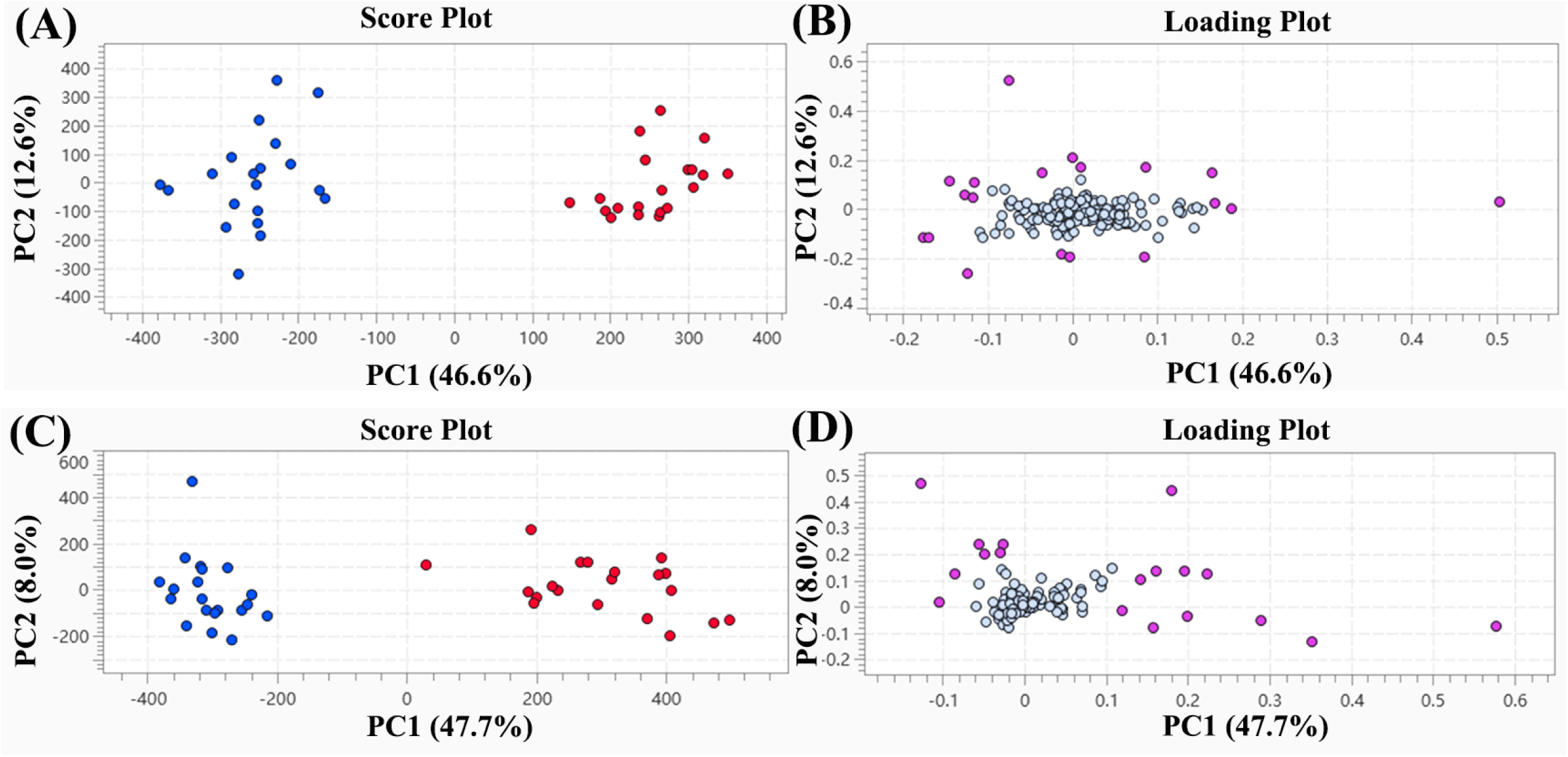
Score plots and loading plots of PCA results. In score plots, red spots represented ROIs selected in non-decayed xylem, while blue spots represented ROIs selected in decayed xylem. Differential markers that contributed most to the difference of decayed and non-decayed xylem were highlighted in the loading plots. (A) Data collected from mass range *m/z* 100-550. (B) Data collected from mass range *m/z* 550-1000.

Ions that contributed most to the chemical difference of decayed and non-decayed xylem of Kuqin were listed in Table 1. Mass intensities of these differential ions were utilized to establish a heatmap (Fig. 6). It was intuitively illustrated by the heatmap that the decayed xylem was much different from the non-decayed xylem. The mass intensities of ions at *m/z* 381.0806, *m/z* 689.2122 and *m/z* 705.1862 were especially high in the non-decayed xylem of Kuqin. Raw data of normalized mass intensities of selected ions could be found in supplemental information (Supplemental Table, S1 to S4).

**Fig. 6.**
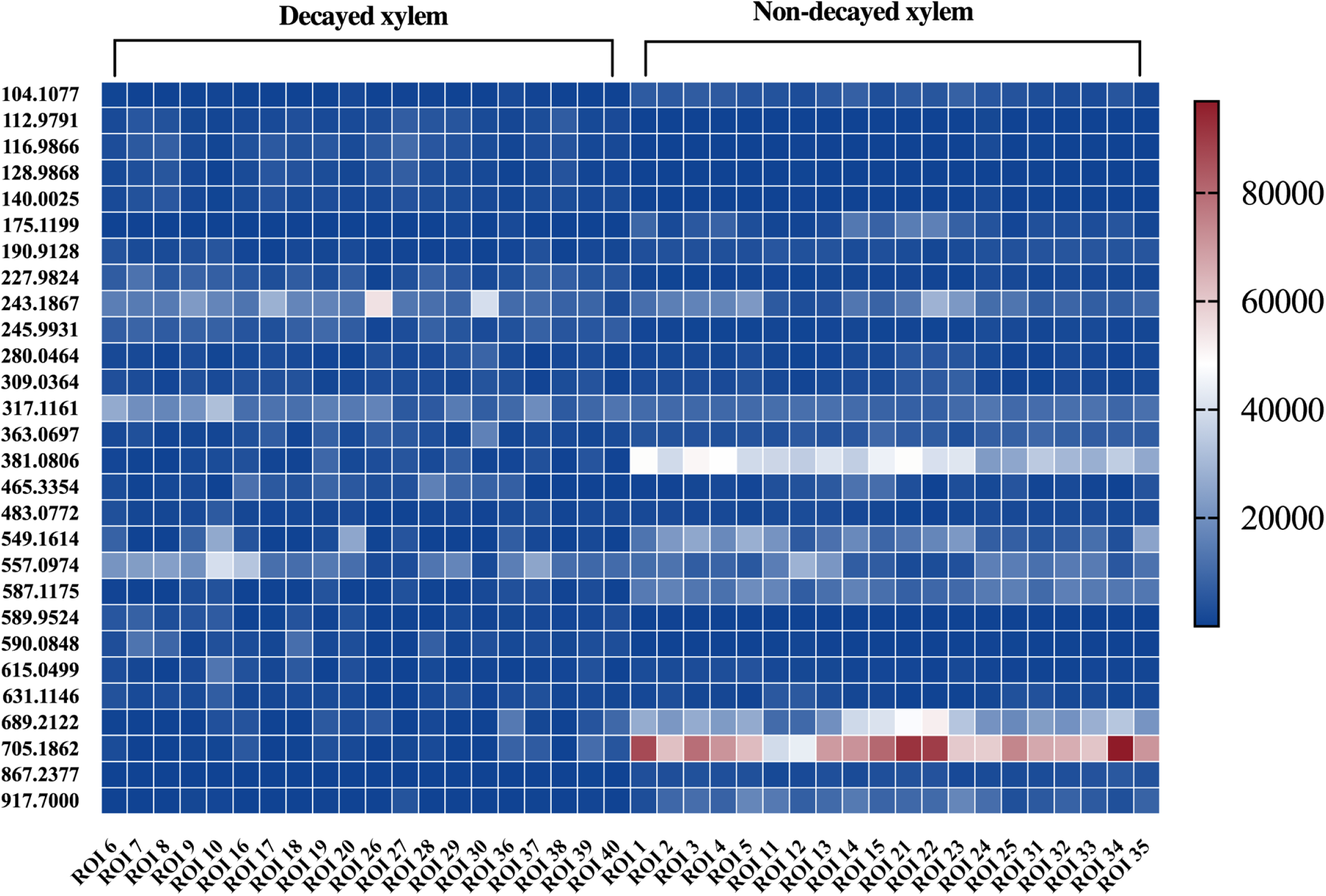
A heatmap established from the raw mass data directly collected from the decayed and non-decayed xylem of Kuqin. The heatmap clearly illustrated the varied mass intensities of differential ions in decayed and non-decayed xylem.

**Table 1.**
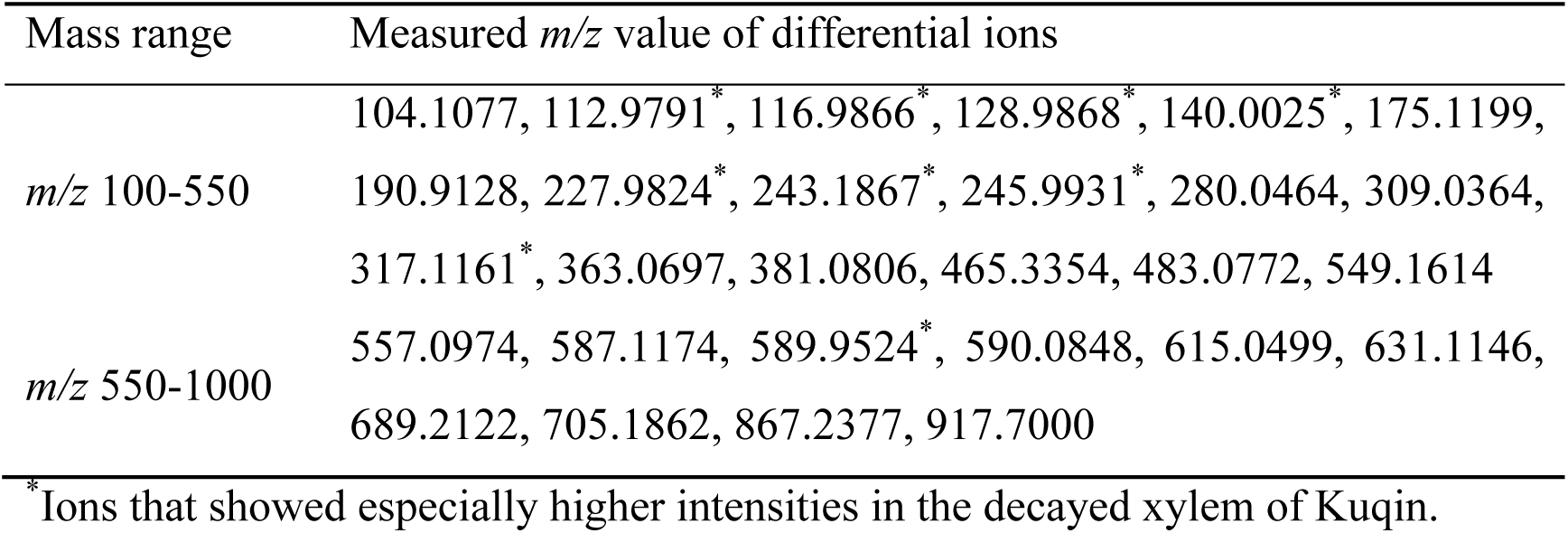
Differential ions of decayed and non-decayed xylem of Kuqin. Ions that presented relatively higher mass intensities in the decayed xylem of Kuqin were marked with asterisks.

| Mass range | Measured $m/z$ value of differential ions |
| --- | --- |
| $m/z$ 100-550 | 104.1077, 112.9791<sup>*</sup>, 116.9866<sup>*</sup>, 128.9868<sup>*</sup>, 140.0025<sup>*</sup>, 175.1199, 190.9128, 227.9824<sup>*</sup>, 243.1867<sup>*</sup>, 245.9931<sup>*</sup>, 280.0464, 309.0364, 317.1161<sup>*</sup>, 363.0697, 381.0806, 465.3354, 483.0772, 549.1614 |
| $m/z$ 550-1000 | 557.0974, 587.1174, 589.9524<sup>*</sup>, 590.0848, 615.0499, 631.1146, 689.2122, 705.1862, 867.2377, 917.7000 |

<sup>\*</sup>Ions that showed especially higher intensities in the decayed xylem of Kuqin.

### Hierarchical clustering analysis summarized the spatial distribution patterns of phytochemicals in Kuqin

While PCA illustrated the chemical difference of decayed and non-decayed xylem from the perspective of spectral difference, hierarchical clustering analysis (HCA) of MS images could explore the chemical difference of two kinds of tissues from the dimension of spatial information. The localization patterns of phytochemicals in Kuqin samples were summarized using HCA. In the dendrogram tree, ions presented similar distribution features were grouped together and occupied the adjacent nodes (Supplemental Fig. S1). Spatial distribution patterns of detected compounds were mainly classified into 4 clusters. Ions of cluster A were found to show especially high abundance in the decayed xylem of Kuqin (Fig. 7A). Ions in cluster B showed preferential distribution in the non-decayed region of Kuqin (Fig. 7B). Ions of cluster C were discovered to exhibit higher intensity in the phloem of samples (Fig. 7C). Ions of cluster D specifically outlined the cork and interxylary cork of samples (Fig. 7D). More information of classification results was supplied in supplemental information (Supplemental Table, S5 and S6).

**Fig. 7.**
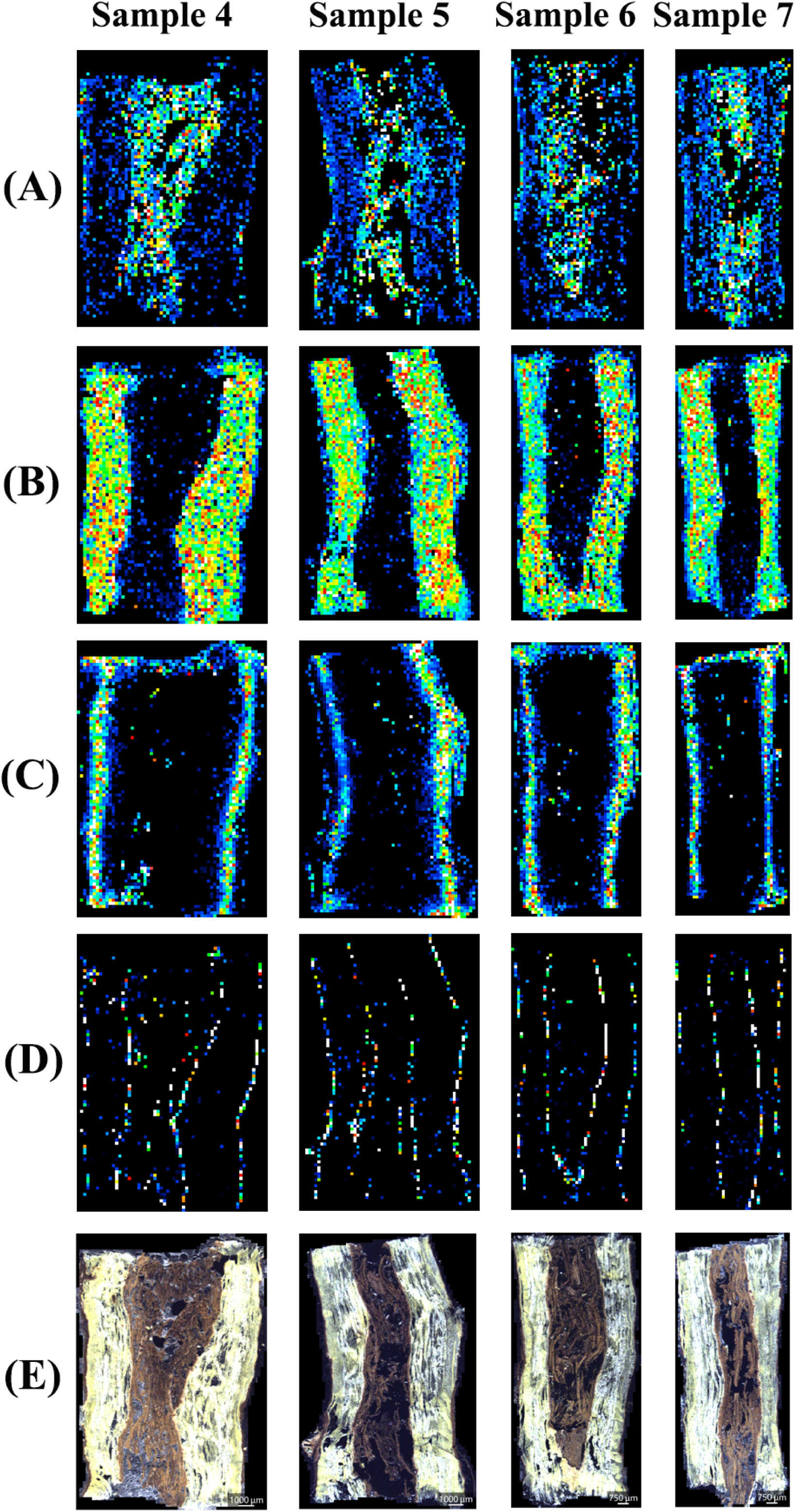
Representative MS images illustrating the spatial distribution of phytochemicals in longitudinal sections of four Kuqin samples. Summarization of the distribution patterns was achieved via HCA. (A) MS images of a representative phytochemical of cluster A, arginine (*m/z* 175.1199, [C_6_H_14_N_4_O_2_+H]^+^, Δ=4.57 ppm). Chemicals of cluster A presented especially high intensities in the decayed xylem. (B) MS images of a representative phytochemical of cluster B, an unknown ion (*m/z* 245.9931). Chemicals of cluster B showed accumulation tendency in non-decayed xylem and non-decayed phloem. (C) MS images of a representative phytochemical of cluster C, sucrose (*m/z* 381.0806, [C_12_H_22_O_11_+H]^+^, Δ=1.84 ppm). Chemicals of cluster C presented especially high intensity in the phloem. (D) MS images of a representative phytochemical of cluster D, skullcapflavone Ⅱ(*m/z* 413.0645, [C_17_H_14_O_8_+K]^+^, Δ=1.45 ppm). This cluster contained only one chemical. (E) Optical images of longitudinal sections of Kuqin samples.

## Discussion

Interxylary cork is an anomalous structure of cork which develops in the secondary xylem of plants. Many phytologists have investigated on the structure and development of interxylary cork using microscope (Fritz & Saukel, 2011; Wang *et al*., 2020). In this experiment, the development of interxylary cork in decayed *S. baicalensis* (Kuqin) was explored using MSI. Spatial distribution of skullcapflavone Ⅱ was regarded as a reflection of interxylary cork development. As a flavanoid aglycone, skullcapflavone Ⅱ is a hydrophobic compound which tends to be soluble in low-polarity solvent. The deposition of suberin in mature cork cell walls might encourage the storage of skullcapflavone Ⅱ (Bernards, 2002), leading to its high mass abundance in cork and interxylary cork regions (Fig. 1B). Similarly, the preferential localization of phytochemicals in cork cells was previously reported in several articles (Pateraki *et al*., 2014; Lange *et al*., 2017). Different from the speculation stated above, these articles hypothesized that the storage of chemicals in cork region was related to the oil bodies that existed in cork cells (Pateraki *et al*., 2014; Lange *et al*., 2017). Both hypothesis emphasized the importance of a lipophilic environment provided by cork cells. In order to determine the specific localization site of lipophilic compounds in cork cells, further work could be carried out by conducting subcellular MSI analysis, which requires higher spatial resolution of MSI equipment. Apart from skullcapflavone Ⅱ, there might also be other compounds that show high accumulation tendency in the suberized cork, however, no compound other than skullcapflavone Ⅱ was found in our experiment. The possible reason might be that the matrix chosen in this experiment was not suitable for the ionization of other lipophilic chemicals. In the future, adjustment of matrices and ionization conditions may be helpful to find out more phytochemicals that showed preferential occurrence in the cork of plants.

The MS images of skullcapflavone Ⅱ intuitively mapped out the successive development of interxylary corks in *S. baicalensis* (Fig. 1B). An unexpected phenomenon observed in our result was the continuous development of interxylary corks within tissues that were already enclosed by an existing interxylary cork, which was not reported in previous investigation on the anatomy of *S. baicalensis* (Wang *et al*., 2020). In an article exploring the development of interxylary cork in *Astragalus membranaceus* (Fisch) Bge. var. *mongholicus* (Bge.) Hsiao, newly formed interxylary corks were only found in the normal xylem tissue. New interxylary corks formed on the outer side of the existing interxylary cork were then connected to the existing one, leading to the expansion of interxylary cork (Han *et al*., 2021). Differently, in another investigation focusing on the anatomical structure of *Gentiana macrophylla* Pall., small-sized interxylary corks were observed in the xylem tissue that was already encircled by a large interxylary cork (Luo, 1987). Our result is consistent with the latter study, proving that multicentric development of small interxylary corks could also be initiated in the abnormal xylem despite its isolation from healthy tissues.

A hypothesis is proposed to explain why small interxylary corks can still be developed within xylem tissues enclosed by an existing interxylary cork. At the beginning of interxylary cork formation, abnormal vessels appear in secondary xylem. Parenchyma cells around these abnormal vessels regain the ability of regeneration (Han *et al*., 2021). Periclinal cell division of dedifferentiated parenchyma cells gives rise to two daughter cells. The smaller daughter cell becomes a cambium cell, while the larger one differentiates into either a cork (phellem) or phelloderm cell (Beck, 2010). As the cork cambium splits in a tangential direction, parenchyma cells which have not triggered regeneration process are enclosed by the newly formed cambium. Afterwards, continuous differentiation of cork cambium produces large quantities of cork cells than phelloderm cells (Beck, 2010), hence the formation of an interxylary cork. Living parenchyma cells which are enclosed by newly formed interxylary cork repeat the process of regeneration and cell division, producing newer interxylary corks within the existing interxylary cork. Therefore, in the pith of sample 1 and sample 2, signals of skullcapflavone Ⅱ were observed in the form of multiple rings (Fig. 1B). Within the large interxylary cork of sample 1, multicentric small interxylary corks were observed. Whereas in sample 2, double layers of interxylary corks presented in the shape of concentric circles. Both anatomical phenomena led to the continuous development of interxylary corks within an existing one. The subsequent formation of interxylary corks is likely to aggravate the isolation of abnormal xylem tissues from healthy tissues, thus accelerates the decaying process of enclosed tissues. Ultimately, all tissues surrounded by the mature interxylary cork are dead. Dead tissues in the decayed pith gradually shed, giving rise to the occurrence of a “hollow heart” in roots.

The successive formation of small interxylary corks in an existing interxylary cork makes no contribution to the expansion of the outermost interxylary cork, but it is closely related to the decaying process of tissues that was enclosed by interxylary corks. As inferred above, tissues surrounded by multiple layers of interxylary cork are isolated from other tissues in an aggravated way. Therefore, we speculate that the existence of successively formed interxylary corks could be utilized to make a segmentation towards tissues that going through different decaying stages. The speculation was further validated by the MS image of baicalein in Kuqin samples (Fig. 2A). An earlier investigation had claimed that baicalein was capable of initiating apoptosis of *Scutellaria* cells (Hirunuma *et al*., 2011). In this experiment, baicalein was observed to show high mass intensity in slightly decayed tissues compared with severely decayed tissues. For instance, in sample 2, the intensity of baicalein in zone 5 was the highest, followed by zone 6 and zone 4 (Fig. 1C, Fig. 2A). In slightly decayed tissue (zone 5), baicalein was produced in a large quantity so as to trigger the programmed cell death. Whereas in non-decayed tissue (zone 6), the mass intensity of baicalein was relatively lower since apoptosis-inducing reagent was not needed in non-decayed tissue. As the decaying process proceeded, baicalein presented a decreased content in severely decayed tissue (zone 4), even lower than the normal tissue, which suggested the possible decomposition of baicalein in dead cells. The spatial distribution of baicalein in sample 1 is very much similar, with a higher mass intensity of baicalein found in the slightly decayed xylem (zone 2) compared with severely decayed xylem (zone 1). Different from the other two samples, in sample 3, no small interxylary cork was newly formed anymore (Fig. 1B), indicating that all cells enclosed by the existing interxylary cork might have lost the ability of regeneration. In other words, the pith of sample 3 decayed in such a severe manner that all cells enclosed by the interxylary cork were probably dead. A very scarce distribution of baicalein was observed in the decayed pith of sample 3 (Fig. 2A), suggesting that baicalein would ultimately be decomposited in the extremely decayed or dead xylem. The spatial distribution of wogonin (Fig. 2B) bore a strong resemblance to baicalein, it could thus be inferred that the physiological function of wogonin might be similar to that of baicalein. Possibly, wogonin was also a compound which could induce cell apoptosis in *S. baicalensis*. In conclusion, the application of MSI not only validated the function of baicalein as an apoptosis-inducing compound in *S. baicalensis*, but also helped to generate new hypothesis of the possible physiological function of wogonin.

Previous studies had demonstrated that the pharmacological effects of decayed and non-decayed roots of *S. baicalensis* differed from each other. One purpose of this study was to uncover the holistic difference of chemical composition between healthy xylem and abnormal xylem of Kuqin. By performing PCA on ROIs drawn in decayed and non-decayed xylem tissues, quite a number of differential ions were sorted out. An interesting aspect of our research was that many differential ions were not reported in previous studies of *S. baicalensis*, thus a majority of differential ions could not be annotated. The possible explanation might be that MSI analysis eliminated the laborious sample preparation procedure, thus avoided the possible loss or degradation of natural compounds that might occur during LC or LC-MS analysis. As a result, compounds which could not been detected by LC or LC-MS were observed by MSI. Therefore, the application of MSI holds potential in revealing new secondary metabolites in plants, which could proceed the progress of phytochemisry. While PCA conducted analysis from the perspective of spectral difference, HCA of MS images of phytochemicals classified detected compounds from the perspective of spatial difference. In HCA results, a large group of phytochemicals from a special cluster were found to show especially high abundance in the decayed pith of roots (Fig. 6A). Since the decayed pith of Kuqin was the representative structure that differed Kuqin from Ziqin, constituents which specifically located here might be the reason why Kuqin held different pharmacological effects from Ziqin. Further research could be undertaken to investigate the bioactivities of differential phytochemicals sorted out by PCA and HCA.

Taken together, this investigation successfully explored the potential of MSI in the exploration of plant science. For the first time, spatial distribution of phytochemicals that stored in certain kinds of tissues was discovered as a reflection of structure development. The serial development of interxylary corks were clearly visualized through the MS image of a special phytochemical. Based on a better comprehension of plant structure, the accumulation tendencies of bioactive components in native tissues were explained in a novel way. Possible physiological function of interest compounds was also speculated according to the spatial information offered by MSI. In the future, the employment of multimodal imaging system is recommended (Buchberger *et al*., 2018). The combination of high-resolution microscopy or histology with MSI may provide supplementary information for the current work.

## Materials and Methods

### Chemicals

2-Nitrophloroglucinol (NPG), acetonitrile, trifluoroacetic acid, 2,5-dihydroxybenoic acid (DHB) and gelatin were purchased from Sigma-Aldrich (St. Louis, MO, United States). Reference standards of baicalein, wogonin, baicalin and wogonosdie were purchased from Shanghai Standard Technology Co. (Shanghai, China). Ultra-pure water was obtained from a Milli-Q water purification system (Millipore, Bedford, MA, United States). Optimum cutting temperature compound (OCT) was purchased from Leica (Nussloch, Germany).

### Plant Materials

Kuqin is the decayed root of *S. baicalensis*. Six Kuqin samples were collected from Neimenggu, China. The origin of samples was authenticated as the root of *S. baicalensis* by Professor Shuai Kang in accordance with the Chinese Pharmacopoeia (Chinese Pharmacopoeia Commission, 2020). For future reference, the voucher specimens were deposited in National Institutes for Food and Drug Control, Beijing, PR China.

### Sample preparation

Kuqin samples were cut into pieces with length of 2-3 cm using a blade. Small pieces of Kuqin were embedded in 0.09 g ml^-1^ gelatin solution respectively. Then the samples were fast-frozen at –80 ℃ until the gelatin turned into solid blocks. Frozen gelatin blocks were axially fixed to a cryomicrotome (Leica, Nussloch, Germany). Roots were transversely or longitudinally sectioned into slices of 30-μm thickness. Root sections were thaw-mounted onto indium tin oxide-coated (ITO) glass slides (Matsunami Glass, Osaka, Japan). Afterwards, matrix deposition was carried out using an automatic sprayer iMLayer AERO (Shimadzu, Tokyo, Japan). NPG solution was prepared at a concentration of 10 mg ml^-1^ in 80% acetonitrile containing 0.1% trifluoroacetic acid. Twenty layers of NPG solution was uniformly deposited onto the root sections.

### Mass spectrometry imaging

DHB solution was dissolved in 50% methanol at a concentration of 10 mg ml^-1^. Mass calibration was performed using DHB solution, the mass error was less than 10 ppm. Prior to matrix deposition, optical images of root sections were captured with a charge coupled device camera attached to the iMScope QT instrument. For the acquisition of mass data, a quadrupole-time-of-flight mass spectrometer equipped with an atmospheric pressure chamber for matrix assisted laser desorption ionization (AP-MALDI) source was utilized. A diode-pumped 355 nm Nd∶YAG laser with a laser repetition frequency of 1000 Hz was utilized for the desorption and ionization of analytes. The diameter of laser beam was set at 25 μm, and the laser intensity was kept at 76.5 (arbitrary unit in iMScope). Root sections were scanned with a raster size of 50 μm. Detection voltage was set at 2.23 kV. The mass spectrometer was operated in positive ion mode. The detection mass range was performed in *m/z* 100-550 and *m/z* 550-1000 for each sample. Mass data of pure NPG was also collected from the ITO glass slides deposited with matrix.

### On-tissue tandem mass spectrometry analysis

On-tissue tandem mass spectrometry was also performed using AP-MALDI-Q-TOF-MS. After matrix application, tandem mass spectra of baicalein, wogonin, baicalin and wogonoside were directly acquired from the transverse sections of Kuqin. Protonated molecules of four flavonoids were selected as the precursor ions. Collision energy was set at 35 ± 15 V. Reference standards of four compounds were dissolved in 70% methanol at a concentration of 10 mg ml^-1^. Standard solution of each flavonoid was dotted onto the surface of a steel plate. Tandem mass spectrometry spectra of reference standards were generated from the dried dots, providing a comparison with the spectra generated from native tissues.

### Data analysis

Prior to further analysis, total ion current (TIC) normalization of mass data was implemented. According to the optical images, regions of interest (ROI) were selected. For each ROI, an average mass spectrum was calculated. By performing peak picking with an intensity threshold of 0.5%, mass peaks exhibiting relatively high mass intensities were included in a data matrix. Detailed information of *m/z* values and corresponding mass intensities were displayed in the data matrix. MS images of different natural compounds were established using ImageReveal MS software.

Twenty ROIs of decayed xylem tissues and twenty ROIs of non-decayed xylem tissues were selected from the longitudinal sections of four Kuqin samples. Mass data generated from two groups of ROIs were utilized to conduct PCA. Compounds having high loading vector were selected as the differential chemicals of decayed and non-decayed xylem of Kuqin. HCA was also employed in this investigation to unveil the different localization tendencies of phytochemicals. Detected compounds were classified into different categories according to the spatial distribution illustrated in MS images.

## Supplemental data

**Supplemental Fig. S1** Dendrogram plot of hierarchical clustering analysis (HCA).

**Supplemental Table S1** Mass intensities of differential ions generated from the average mass spectra of Kuqin (Sample 4).

**Supplemental Table S2** Mass intensities of differential ions generated from the average mass spectra of Kuqin (Sample 5).

**Supplemental Table S3** Mass intensities of differential ions generated from the average mass spectra of Kuqin (Sample 6).

**Supplemental Table S4** Mass intensities of differential ions generated from the average mass spectra of Kuqin (Sample 7).

**Supplemental Table S5** Classification of ions presenting different distribution patterns (mass range *m/z* 100-550).

**Supplemental Table S6** Classification of ions presenting different distribution patterns (mass range *m/z* 550-1000).

## Acknowledgements and funding

This work was supported by the National Natural Science Foundation of China (81973476) and ASEAN-China Coorperation Fund (ACCF) (Ref. No. ASCC/HDD/HLD/LET/2023/29).

## Author contributions

L. H. and L. N. coordinated the research and wrote the manuscript. L. H. and L.N. designed and performed the research. J. D. assisted with MSI operation. L.Y. assisted with data analysis. S. K. assisted with sample collection and sample preparation. H. J., F. W. and S. M. revised the manuscript. L.H. and L. N. contributed equally.

